# Spatiotemporal binding of cyclophilin A and CPSF6 to capsid regulates HIV-1 nuclear entry and integration

**DOI:** 10.1101/2024.04.08.588584

**Authors:** Zachary Ingram, Christopher Kline, Alexandra K. Hughson, Parmit K. Singh, Hannah L. Fischer, Gregory A. Sowd, Simon C. Watkins, Melissa Kane, Alan N. Engelman, Zandrea Ambrose

## Abstract

Human immunodeficiency virus type 1 (HIV-1) capsid, which is the target of the antiviral lenacapavir, protects the viral genome and binds multiple host proteins to influence intracellular trafficking, nuclear import, and integration. Previously, we showed that capsid binding to cleavage and polyadenylation specificity factor 6 (CPSF6) in the cytoplasm is competitively inhibited by cyclophilin A (CypA) binding and regulates capsid trafficking, nuclear import, and infection. Here we determined that a capsid mutant with increased CypA binding affinity had significantly reduced nuclear entry and mislocalized integration. However, disruption of CypA binding to the mutant capsid restored nuclear entry, integration, and infection in a CPSF6-dependent manner. Furthermore, relocalization of CypA expression from the cell cytoplasm to the nucleus failed to restore mutant HIV-1 infection. Our results clarify that sequential binding of CypA and CPSF6 to HIV-1 capsid is required for optimal nuclear entry and integration targeting, informing antiretroviral therapies that contain lenacapavir.

## Introduction

The human immunodeficiency virus type 1 (HIV-1) capsid is a unique structure formed from capsid protein (CA) monomers that assemble into approximately 250 hexamers and 12 pentamers (Ganser et al., 1999; Zhao et al., 2013). Following HIV-1 fusion with the target cell membrane, the capsid enters the cytoplasm where it protects the two copies of viral RNA from recognition by host innate immune factors (Yamashita and Engelman, 2017). Simultaneously, the HIV-1 capsid surface is an interface for binding host motor proteins and their adapters to promote directed microtubule trafficking (Lukic et al., 2014; Malikov et al., 2015; Stephens and Naghavi, 2022). At the nucleus, specific host nucleoporin proteins bind to HIV-1 capsid to facilitate nuclear import of the HIV-1 genome (Di Nunzio et al., 2012; Dickson et al., 2024; Fu et al., 2024; Matreyek et al., 2013; Shen et al., 2023a; Shen et al., 2023b).

Throughout these post-entry steps, HIV-1 capsid undergoes a poorly defined disassembly process, termed capsid uncoating, which is affected by host proteins (Ambrose and Aiken, 2014; Campbell and Hope, 2015). Capsid uncoating likely occurs at the nuclear pore complex (NPC) or within the nucleus, allowing completion of reverse transcription of the viral genome into double-stranded DNA, which is integrated into host chromatin (Burdick et al., 2017; Burdick et al., 2020; Dharan et al., 2020; Muller et al., 2021; Zila et al., 2021). As HIV-1 capsid has a unique structure and is involved in numerous post-entry steps, it is a useful target for development of antiretroviral therapeutics. Lenacapvir (LCV) is the first antiretroviral inhibitor that targets HIV-1 capsid that has been approved for use in humans (Link et al., 2020; Segal-Maurer et al., 2022). Interestingly, LCV inhibits multiple steps of HIV-1 replication (Bester et al., 2020; Link et al., 2020).

Optimal HIV-1 replication depends on capsid binding to multiple host proteins. Cyclophilin A (CypA) is an abundant cellular peptidyl prolyl isomerase involved in protein folding and trafficking (Koletsky et al., 1986; McCaffrey et al., 1993; Zhang et al., 2013) and binds to the partially ordered loop between alpha helices 4 and 5 of HIV-1 CA (Gamble et al., 1996; Luban et al., 1993). CypA promotes early HIV-1 replication steps in a cell type dependent manner that correlates with events prior to nuclear import (Braaten et al., 1996; De Iaco and Luban, 2014; Sokolskaja et al., 2004). Recently it was reported that CypA binding to HIV-1 capsid prevents binding of the restriction factor TRIM5_α_ to capsid in primary human CD4+ T cells and macrophages (Kim et al., 2019; Selyutina et al., 2020). TRIM5_α_ is an antiretroviral restriction factor that oligomerizes on the HIV-1 capsid surface, forming hexagonal nets that destabilize capsid and impair HIV-1 replication (Black and Aiken, 2010; Li et al., 2016; Stremlau et al., 2004). Beyond establishing a mechanism for how CypA promotes HIV-1 replication, these findings suggest that host factors compete for binding to the HIV-1 capsid surface and can alter replication outcomes.

Similarly, our group previously showed that CypA binding limits HIV-1 capsid binding to the host protein cleavage and polyadenylation specificity factor 6 (CPSF6) in the cytoplasm, impacting capsid uncoating, nuclear trafficking, and infection (Ning et al., 2018; Zhong et al., 2021). CPSF6 is localized to the nucleus through binding of its C-terminal RS-like domain to transportin 3 (TNPO3) (Jang et al., 2019; Lee et al., 2010; Maertens et al., 2014), where it functions as a pre-mRNA alternate polyadenylation factor (Dettwiler et al., 2004). CPSF6 directly binds to HIV-1 capsid within a hydrophobic pocket between adjacent HIV-1 CA monomers (Bhattacharya et al., 2014; Lee et al., 2010; Price et al., 2012), which is also the binding site of LCV (Bester et al., 2020). CPSF6 promotes HIV-1 capsid disassembly, nuclear import, and genome integration within gene dense chromatin and speckle-associated domains (SPADs) (De Iaco et al., 2013; Francis et al., 2020b; Zhong et al., 2021).

Research on individual host interactions with HIV-1 capsid has elucidated key HIV-1 replication events. Here, we have explored the spatiotemporal kinetics of CypA and CPSF6 binding to HIV-1 capsid and the impact these interactions have on HIV-1 replication, using both CA mutants and inhibitors. The unique mutant HIV-1_AC-1_ was shown to have increased CypA binding affinity, resulting in failed nuclear import in all cell types tested unless CypA binding to capsid was inhibited (Lahaye et al., 2013; Lahaye et al., 2016). Introduction of the CA amino acid substitution N74D reversed the replication defect of HIV-1_AC-1_ by an undetermined mechanism (Lahaye et al., 2016). N74D was shown to abolish CPSF6 binding, leading to HIV-1 infection via alternative NPC components and genome integration outside of transcriptionally active genes (Lee et al., 2010; Sowd et al., 2016). Given an incomplete understanding of HIV-1_AC-1_ restriction and our previously described relationship of CypA and CPSF6 binding to capsid, we hypothesized that increased CypA binding affinity to capsid prevents CPSF6 from accessing the HIV-1 capsid, leading to reduced nuclear import, integration, and infection. Loss of CypA binding would restore HIV-1_AC-1_ capsid binding to CPSF6 and infection, suggesting sequential binding of host factors to capsid is required for replication.

## Results

### Increased CypA binding inhibits HIV-1 infection in a CPSF6-dependent manner

The HIV-1_AC-1_ mutant was engineered by introduction of five amino acid substitutions within the CypA binding loop of CA, V86I, I91L, A92P, P93A and M96L (**Figure 1A**), resulting in increased CypA binding affinity and significant inhibition of infection (Lahaye et al., 2013). Infectivity of HIV-1_AC-1_ was rescued to near wild-type (WT) levels by treatment of the cells with cyclosporine A (CsA), which is a calcineurin inhibitor that prevents CypA from binding to HIV-1 capsid, or by reducing CypA expression in target cells (Lahaye et al., 2016). To validate the previously reported phenotype, we compared WT HIV-1 and HIV-1_AC-1_ infection in different cell types with different methods of inhibiting CypA binding to capsid. In agreement with the literature, HIV-1_AC-1_ infection was significantly restricted in HeLa (**Figure 1B**), Jurkat (**Figure 1C**), and primary CD4+ T (**Figure 1D**) cells and these defects were rescued to WT levels in the presence of CsA (WT + CsA). In addition, we compared WT HIV-1 and HIV-1_AC-1_ infectivity in Jurkat T cells lacking the *PPIA* gene, which encodes CypA (Sokolskaja et al., 2004). In *PPIA^-/-^* cells, WT HIV-1 and HIV-1_AC-1_ supported similar levels of infectivity in the presence or absence of CsA (**Figure 1E**). Importantly, HIV-1_AC-1_ had similar infectivity as WT HIV-1 in *PPIA^-/-^* cells, confirming that CypA binding leads to HIV-1_AC-1_ restriction. This was further confirmed by addition of the P90A CA mutation, which prevents CypA binding to HIV-1 capsid (Franke et al., 1994; Wiegers et al., 1999), in HIV-1_AC-1_, resulting in infectivity comparable to P90A HIV-1, regardless of CsA treatment (**Figure S1A**).

**Figure 1:**
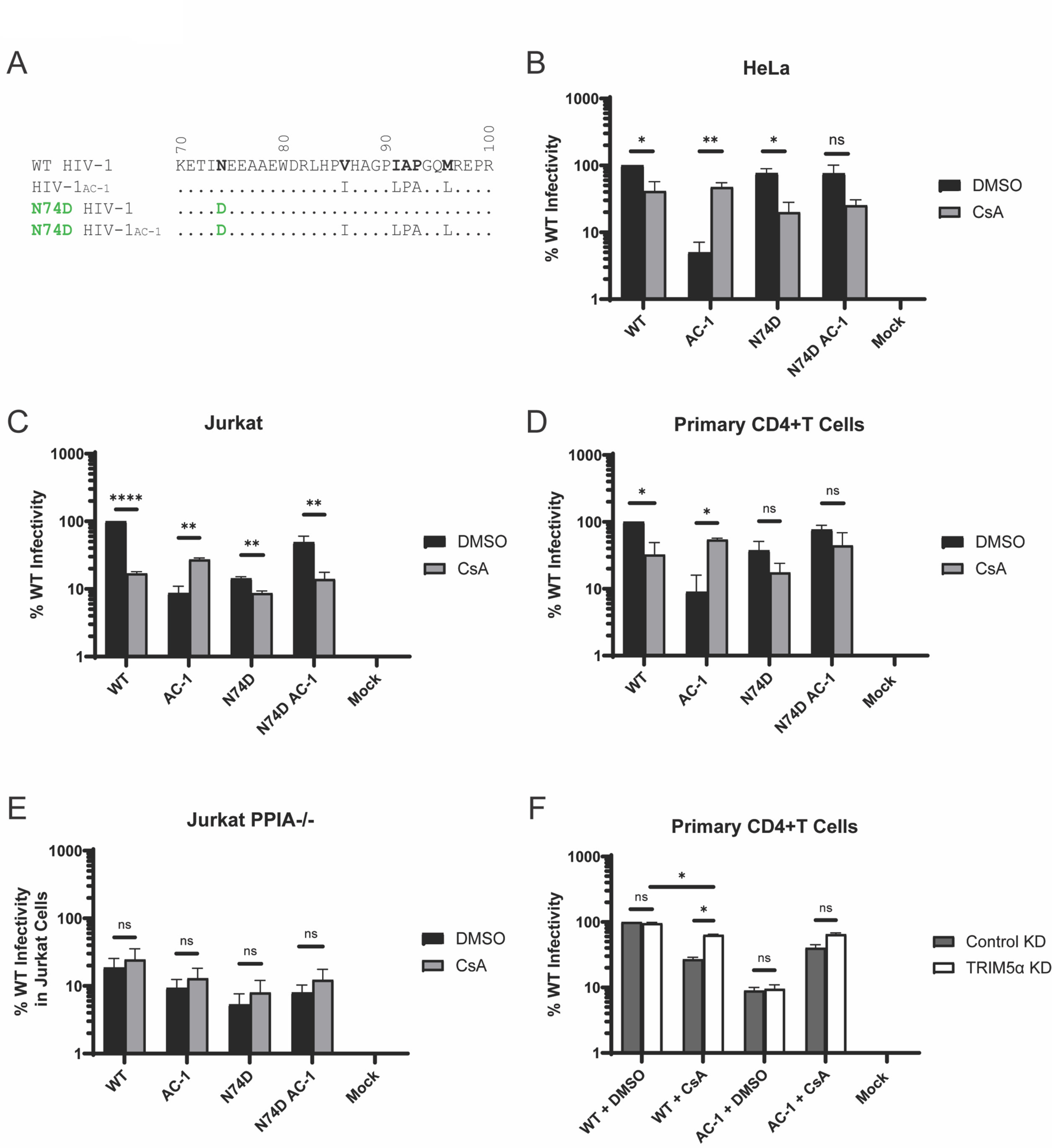
Increased CypA binding to HIV-1 capsid inhibits infection in a CPSF6-dependent manner. (A) CA sequences (amino acids 70-100) are compared between WT HIV-1 and mutants. (B-E) Infectivity of WT and mutant HIV-1 (10 ng p24) was determined after 48 h by luciferase activity in DMSO or 5-10 μM CsA in HeLa (n = 3; B), Jurkat (n = 3; C), Jurkat *PPIA^-/-^* (n = 3; D), and primary CD4+ T cells (n = 2; E). (F) Primary CD4+ T cells were transduced with lentiviruses expressing control or TRIM5_α_ miRNA prior to infection with WT and mutant HIV-1 (10 ng p24) in DMSO or 10 μM CsA. Infections were determined after 48 h by luciferase activity (n = 2). Error bars represent standard errors of the mean (SEM). Comparisons between infection conditions were analyzed by unpaired t tests. P values of < 0.05 were considered significant and significant values are denoted as *, p < 0.05; **, p < 0.01; ***, p < 0.001; and ****, p < 0.0001. ns, p > 0.05.

The CA N74D substitution abolishes HIV-1 capsid binding to CPSF6 but maintains CypA binding (Ambrose et al., 2012; Lee et al., 2010). Addition of N74D to HIV-1_AC-1_ was shown to reverse the infectivity restriction of HIV-1_AC-1_ (Lahaye et al., 2016). Indeed, we show no difference in infection of N74D HIV-1 and N74D HIV-1_AC-1_ in HeLa, Jurkat T, and primary CD4+ T cells (**Figures 1B, 1C, 1D, 1E**). These results suggested that CPSF6 binding also is required for HIV-1_AC-1_ restriction.

As previous work suggested that CypA binding to HIV-1 capsid shields it from TRIM5_α_ binding in human primary cells (Kim et al., 2019; Selyutina et al., 2020), we hypothesized that HIV-1_AC-1_ also would be resistant to TRIM5_α_ restriction. Primary CD4+ T cells were transduced with lentiviral vectors expressing miRNA targeting TRIM5_α_ or a control miRNA. TRIM5_α_ knockdown (KD) was confirmed by infection with N-tropic murine leukemia virus (N-MLV), which is restricted by TRIM5_α_ (Pertel et al., 2011; Yap et al., 2004) (**Figure S1B**). As previously reported, WT HIV-1 infection was inhibited by CsA treatment, which was partially restored by TRIM5_α_ KD (**Figure 1F**). As expected, HIV-1_AC-1_ infectivity was not further restricted by TRIM5_α_ KD in the absence of CsA. However, infectivity of HIV-1_AC-1_ was increased by TRIM5_α_ KD in the presence of CsA, but to a lesser extent than WT HIV-1, which could be due to the higher CypA binding affinity. Thus, CypA binding to HIV-1_AC-1_ capsid prevents TRIM5_α_ restriction like WT capsid in primary CD4+ T cells. But this does not account for the general restriction of HIV-1_AC-1_ in both primary cells and cell lines.

### Increased CypA binding did not affect inherent HIV-1 capsid stability

To characterize the relative stabilities of WT HIV-1 and HIV-1_AC-1_ capsids, we used the *in vitro* CA retention assay (Xu et al., 2020). After adhering to glass and disrupting the envelope, virions containing fluorescently tagged integrase (IN) were fixed, stained with an anti-CA antibody, and imaged by microscopy (**Figure 2A**). CA retention of HIV-1_AC-1_ was comparable to WT HIV-1 CA (∼50% of initial CA signal), suggesting that the inherent stabilities of the respective viral capsids are similar (**Figures 2B, 2C**). The HIV-1 CA mutants K203A and E45A were included as hypostable and hyperstable controls (Forshey et al., 2002), which display rapid loss of CA and delayed retention of CA, respectively (Xu et al., 2020). As expected, K203A capsids showed a significant decrease in retained CA (< 20%), whereas E45A capsids maintained nearly 90% of the initial CA staining signal (**Figure 2C**).

**Figure 2:**
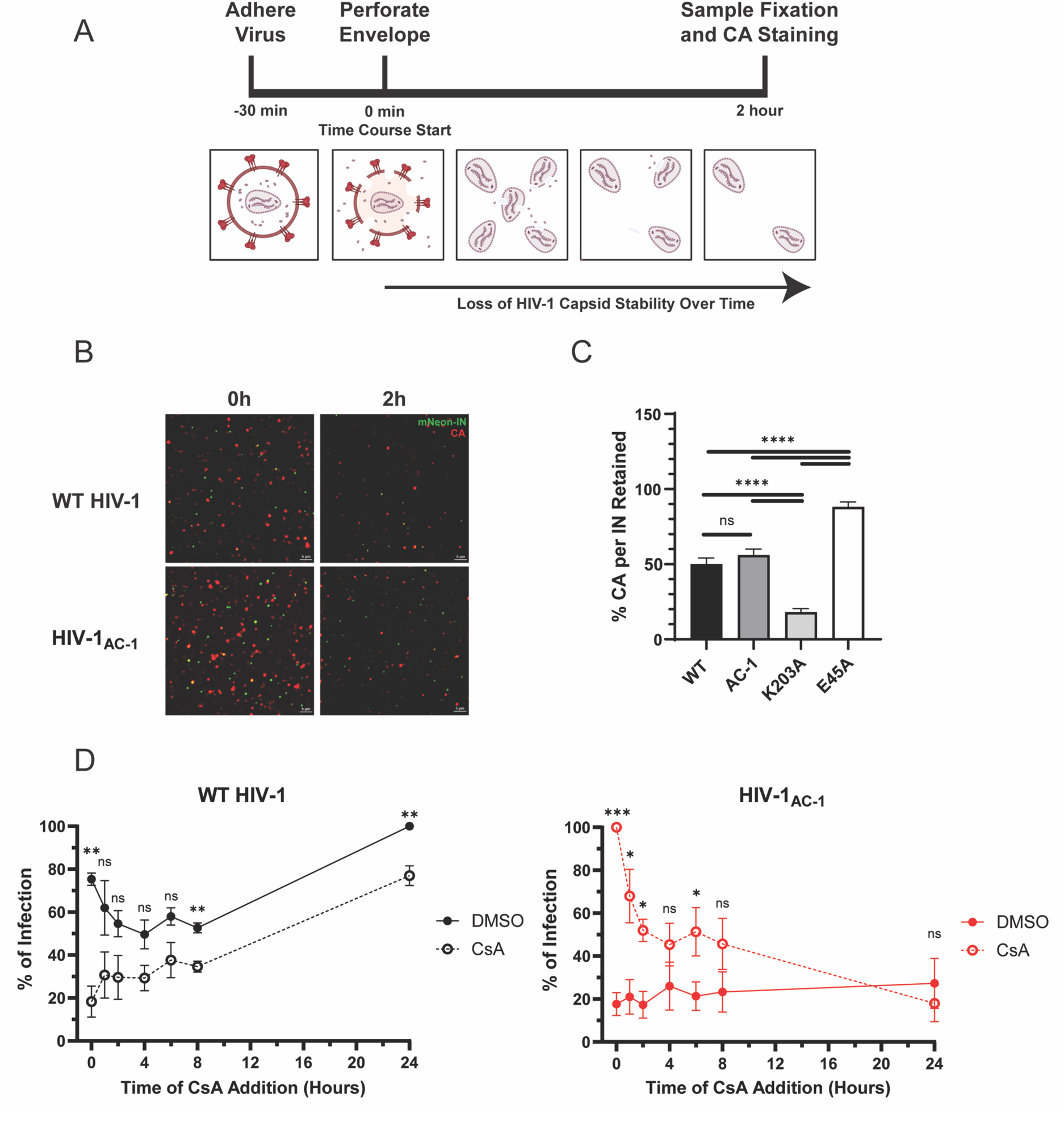
HIV-1_AC-1_ has similar capsid stability to WT HIV-1. (A) Schematic of the HIV-1 CA retention assay. (B) Representative TIRF images of HIV-1 samples fixed at 0 h and 2 h are shown. Scale bars denote 5 μm. (C) Quantification of the average HIV-1 CA retention normalized to mRuby3-IN for WT HIV-1 and HIV-1_AC-1_ (n = 3, 8 fields/virus). (D) DMSO or 10 μM CsA was added at different time points to HeLa cells synchronously infected with WT HIV-1 or HIV-1_AC-1_ (n = 3). Infectivity was normalized to maximal infection (24 h in DMSO for WT, 0 h in CsA for AC-1). Error bars represent SEM and unpaired t-tests were performed for comparisons between viruses (C) or conditions at each time point (D). P values < 0.05 were considered significant and significant values are denoted as *, p < 0.05; **, p < 0.01; ***, p < 0.001; and ****, p < 0.0001. ns, p > 0.05.

Given similar *in vitro* stability of HIV-1_AC-1_ and WT HIV-1 capsids, we investigated whether HIV-1_AC-1_ capsid is stable in cells and whether binding to CypA leads to irreversible restriction of infection. HeLa cells were synchronously infected with WT HIV-1 or HIV-1_AC-1_. CsA-containing medium was introduced at different time points to disrupt CypA binding. Infection was measured 48 h later to assess at which time points CsA could still rescue HIV-1_AC-1_ infectivity (**Figures 2D, S1C, S1D**). Medium containing the same concentration of DMSO was used as a control. CsA addition to WT HIV-1 at the onset of infection resulted in a significant drop in infectivity, which is likely due to CypA enhancing early replication events (De Iaco and Luban, 2014). In contrast, HIV-1_AC-1_ infectivity was maximal when CsA was introduced immediately during infection and then decreased over time. However, HIV-1_AC-1_ infection could be partially rescued by CsA up to 6 h post-infection, after which the addition of CsA did not significantly restore infectivity, indicating that HIV-1_AC-1_ capsid was not immediately destabilized by CypA binding but experienced an irreversible change by 6 h after infection.

### Increased CypA binding to HIV-1 capsid leads to failed nuclear import

As HIV-1_AC-1_ capsid stability is comparable to WT HIV-1, we investigated the reported defect in completing nuclear import (Lahaye et al., 2016). To determine if CypA binding to HIV-1_AC-1_ impacted reverse transcription, WT HIV-1 and HIV-1_AC-1_ *gag* DNA copies were quantified 24 h post-infection (**Figure 3A**). HIV-1_AC-1_ had no difference in the number of reverse transcripts compared to WT HIV-1, regardless of CsA treatment, suggesting that reverse transcription is not impaired. Nuclear import was measured by quantitation of HIV-1 2-long terminal repeat (LTR)-containing circles, which form by non-homologous end joining of unintegrated HIV-1 DNA in the nucleus (Li et al., 2001) (**Figure 3B**). HIV-1_AC-1_ infection led to lower levels of 2-LTR circles, which were restored to WT HIV-1 levels when the cells were treated with CsA.

**Figure 3:**
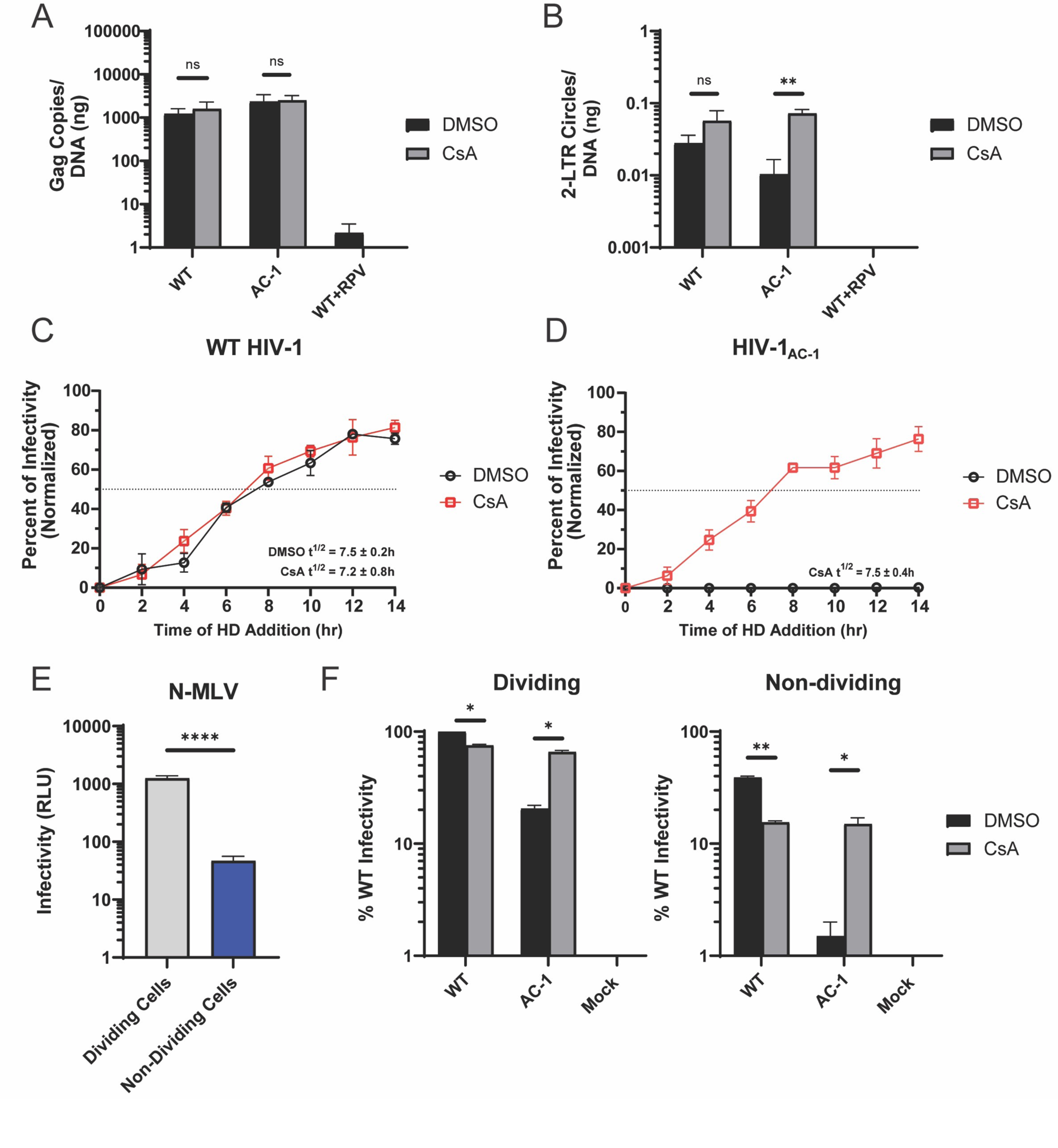
Increased CypA binding restricts HIV-1 nuclear import. (A) Reverse transcripts and (B) 2-LTR circles were measured by qPCR for WT HIV-1 and HIV-1_AC-1_ infection of HeLa cells treated with DMSO or 10 μM CsA after 24 h (n = 3). (C-D) Nuclear import kinetics were determined for WT HIV-1 (C) and HIV-1_AC-1_ (D) infection of HeLa cells in DMSO or 10 μM CsA containing media at different time points (n = 3). Infectivity was determined by luciferase activity and normalized to luciferase expression at 24 h post-infection. Horizontal dotted lines intersect the y-axis at 50% infectivity. (E-F) HeLa cells treated with or without aphidicolin were infected with N-MLV encoding luciferase (E) or WT HIV-1 and HIV-1_AC-1_ encoding mNeonGreen (F) in the presence of DMSO or 10 μM CsA (n = 2). Infectivity was measured by reporter gene expression at 48 h. Error bars represent SEM. Comparisons between infection conditions were analyzed by unpaired t tests. P values < 0.05 were considered significant and significant values are denoted as *, p < 0.05; **, p < 0.01; ***, p < 0.001; and ****, p < 0.0001. ns, p > 0.05.

While 2-LTR circle quantitation is common for assessing HIV-1 nuclear import, they form after nuclear import and require completion of reverse transcription. Thus, the detection of 2-LTR circles may not accurately reflect the kinetics of nuclear import. Thus, the nuclear import kinetics (NIK) assay was performed, in which expression in cells of a fusion Nup62-GFP containing a drug inducible dimerizing domain leads to blockage of the central channel of the NPC when a homodimerization (HD) drug is introduced (Dharan et al., 2020). Addition of HD drug at different time points prevents nuclear import of HIV-1 genomes, corresponding to a decrease in infectivity over time. The NIK assay was performed in cells infected with WT HIV-1 or HIV-1_AC-1_ in the presence or absence of CsA. The WT HIV-1 infection half-life in the NIK assay was calculated as 7.5± 0.2 h post-infection without CsA and 7.1 ± 0.8h with CsA (**Figures 3C, S2A**), which are consistent with a previous report (Dharan et al., 2020). In contrast, HIV-1_AC-1_ nuclear import was undetectable in the presence of DMSO over the 14 h experimental time course, further indicating that this mutant is restricted at nuclear import (**Figures 3D, S2B**). However, HIV-1_AC-1_ nuclear import in the presence of CsA resulted in an infection half-life of 7.5 ± 0.4 h, suggesting that HIV-1_AC-1_ nuclear import proceeds similarly to that of WT HIV-1 only when CypA binding is disrupted.

As the HIV-1 genome can access the nucleus via both import through the NPC and breakdown of the nuclear envelope during mitosis, we hypothesized that restriction of HIV-1_AC-1_ infection would be enhanced in nondividing cells. HeLa cells were infected with viruses in the presence or absence of aphidicolin. Infection of aphidicolin-treated cells with N-MLV, which cannot infect nondividing cells (Lewis et al., 1992; Roe et al., 1993), served as a control for aphidicolin treatment (**Figure 3E**). Infection of WT HIV-1 was reduced in nondividing cells in both the presence and absence of CsA treatment (**Figure 3F**). In contrast, HIV-1_AC-1_ infectivity was more dramatically reduced in nondividing cells in DMSO. Addition of CsA rescued HIV-1_AC-1_ infectivity to WT HIV-1 + CsA levels, suggesting that HIV-1_AC-1_ can use NPCs in nondividing cells when CypA binding is inhibited.

### Cytoplasmic but not nuclear CypA regulates HIV-1 nuclear import and infection

We previously showed in HeLa and SupT1 cells that CypA expression is predominantly cytoplasmic with no localization in the perinuclear region overlapping with the microtubule organizing center or in the nucleus (Zhong et al., 2021). We proposed that cytoplasmic CypA prevents premature cytoplasmic CPSF6 binding to HIV-1 capsid during trafficking towards the nucleus. Staining of CypA in HT1080 cells showed a similar expression pattern (**Figure 4A**).

**Figure 4:**
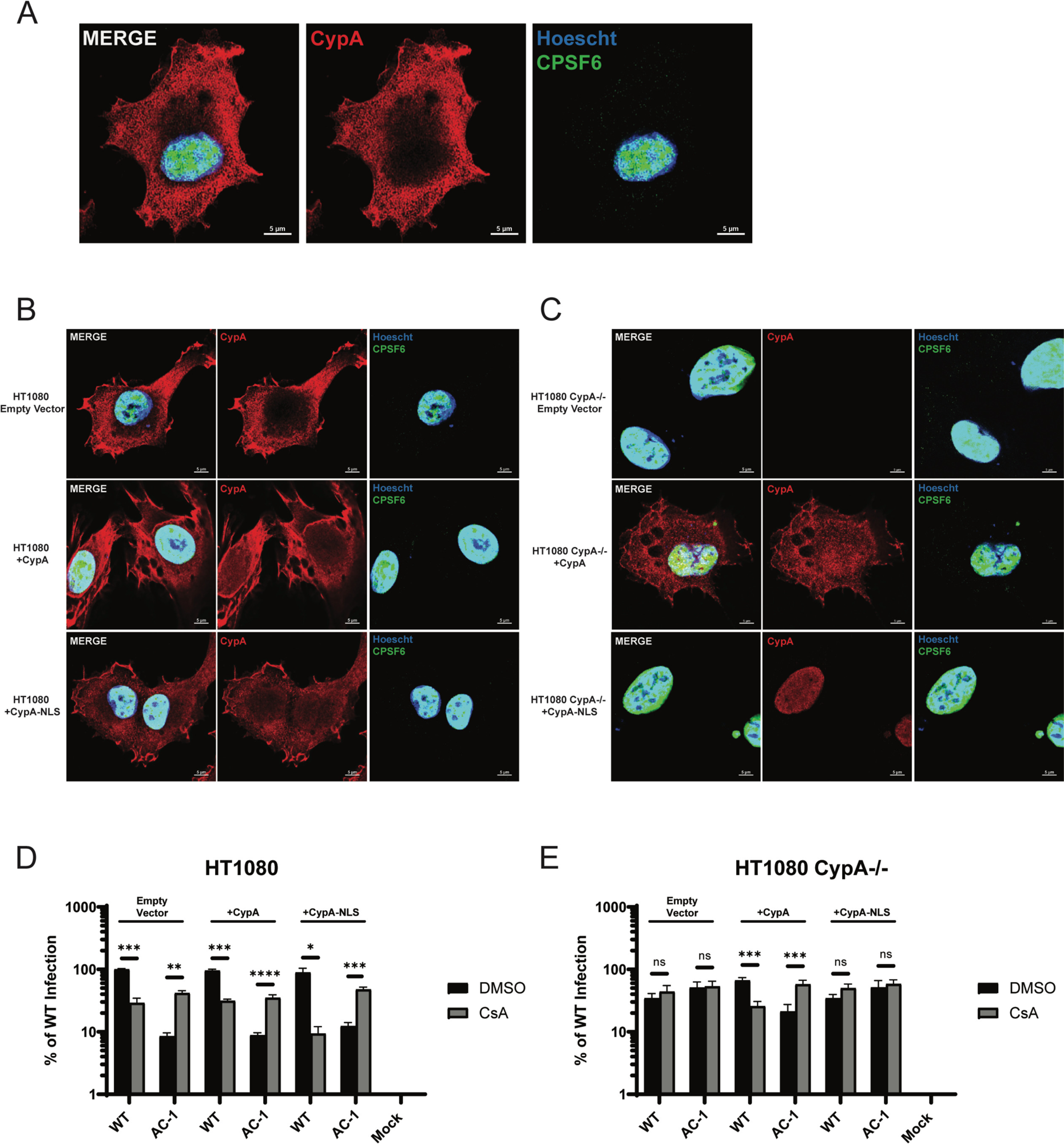
Cytoplasmic but not nuclear CypA regulates HIV-1 nuclear import and infection. (A) Representative confocal microscopy image of HT1080 cells stained with Hoescht (blue) and antibodies against CypA (red) and CPSF6 (green). (B-C) Representative images of HT1080 control cells (B) or HT1080 CypA^-/-^ cells (C) transfected with an empty control plasmid or a plasmid encoding either CypA or CypA-NLS and stained for CypA, CPSF6, and nuclei. (D-E) Infection of transfected HT1080 control cells (D) or HT1080 CypA^-/-^ cells (E) was measured at 48 h in media containing DMSO or 10 μM CsA (n = 2). Error bars represent SEM. Comparisons between infection conditions were analyzed by unpaired t tests. P values < 0.05 were considered significant and significant values are denoted as *, p < 0.05; **, p < 0.01; ***, p < 0.001; and ****, p < 0.0001. ns, p > 0.05.

As increased CypA binding results in reduced HIV-1_AC-1_ nuclear import and infection, we sought to address whether cytoplasmic localization of CypA is required for nuclear import. To test this, we used a HT1080 cell line that was CRISPR-edited to delete the *PPIA* gene (Layish et al., 2024). Transfection of a plasmid encoding CypA into HT1080 control and CypA^-/-^ cells led to CypA expression throughout the cell (**Figures 4B, 4C**). In contrast, addition of the SV40 nuclear localization signal to CypA (CypA-NLS) led to exclusive nuclear expression in HT1080 CypA^-/-^ cells (**Figure 4C**). WT HIV-1 and HIV-1_AC-1_ similarly infected control HT1080 cells regardless of CsA treatment or after transfection with empty vector or vectors expressing CypA or CypA-NLS (**Figure 4D**), which was consistent with other cell types (**Figures 1B, 1C, 1E**). In HT1080 CypA^-^/^-^ cells, HIV-1_AC-1_ infection was rescued to WT HIV-1 levels regardless of CsA addition (**Figure 4E**), which was similar to what was observed in Jurkat *PPIA*^-^/^-^ cells (**Figure 1D**). While exogenous expression of CypA in HT1080 CypA^-/-^ cells reverted both WT HIV-1 and HIV-1_AC-1_ infection patterns to that of control HT1080 cells, CypA-NLS transfection did not (**Figure 4E**). These results suggest that CypA expression in the cytoplasm of cells regulates nuclear entry and infection.

### Increased CypA binding to HIV-1 capsid leads to reduced CPSF6 and LCV binding

Given the rescue of HIV-1_AC-1_ infection by addition of the N74D substitution and that CypA binding limits CPSF6 binding to WT HIV-1 capsid, we investigated the interaction between HIV-1_AC-1_ capsid and CPSF6. We first addressed whether HIV-1_AC-1_ can engage nuclear CPSF6 during infection. Following WT HIV-1 nuclear entry, CPSF6 tagged with a fluorophore can form visible higher order complexes, suggesting binding to capsid (Francis et al., 2020b; Xue et al., 2023). HeLa cells expressing GFP-CPSF6 were synchronously infected with equivalent numbers of WT HIV-1 or HIV-1_AC-1_ particles in the presence or absence of CsA for 6 h before fixation and imaging. WT HIV-1 infection resulted in visible CPSF6 higher order complexes with or without CsA treatment, with cells averaging 6 or 8 per nuclei, respectively (**Figures 5A, 5B**). In contrast, few CPSF6 complexes were detected after HIV-1_AC-1_ infection unless CsA was added (**Figure 5A**). Quantification of the higher order complexes in the absence of CsA treatment showed an average of 0.9 CPSF6 complexes per nuclei (**Figure 5B**), likely due to the lack of nuclear entry when CypA is bound to HIV-1_AC-1_ capsid. Addition of CsA restored formation of CPSF6 complexes by HIV-1_AC-1_ infection to an average of 8 CPSF6 complexes per nuclei, comparable to WT HIV-1 levels. As a control, HeLa cells expressing GFP-CPSF6 with the F284A substitution, which prevents HIV-1 capsid binding (Francis et al., 2020b; Sowd et al., 2016), were infected with WT HIV-1. Only 1.5 F284A CPSF6 higher order complexes per nuclei were detected, indicating that the puncta were specific to CPSF6 interaction with HIV-1 capsid (**Figure S3**).

**Figure 5:**
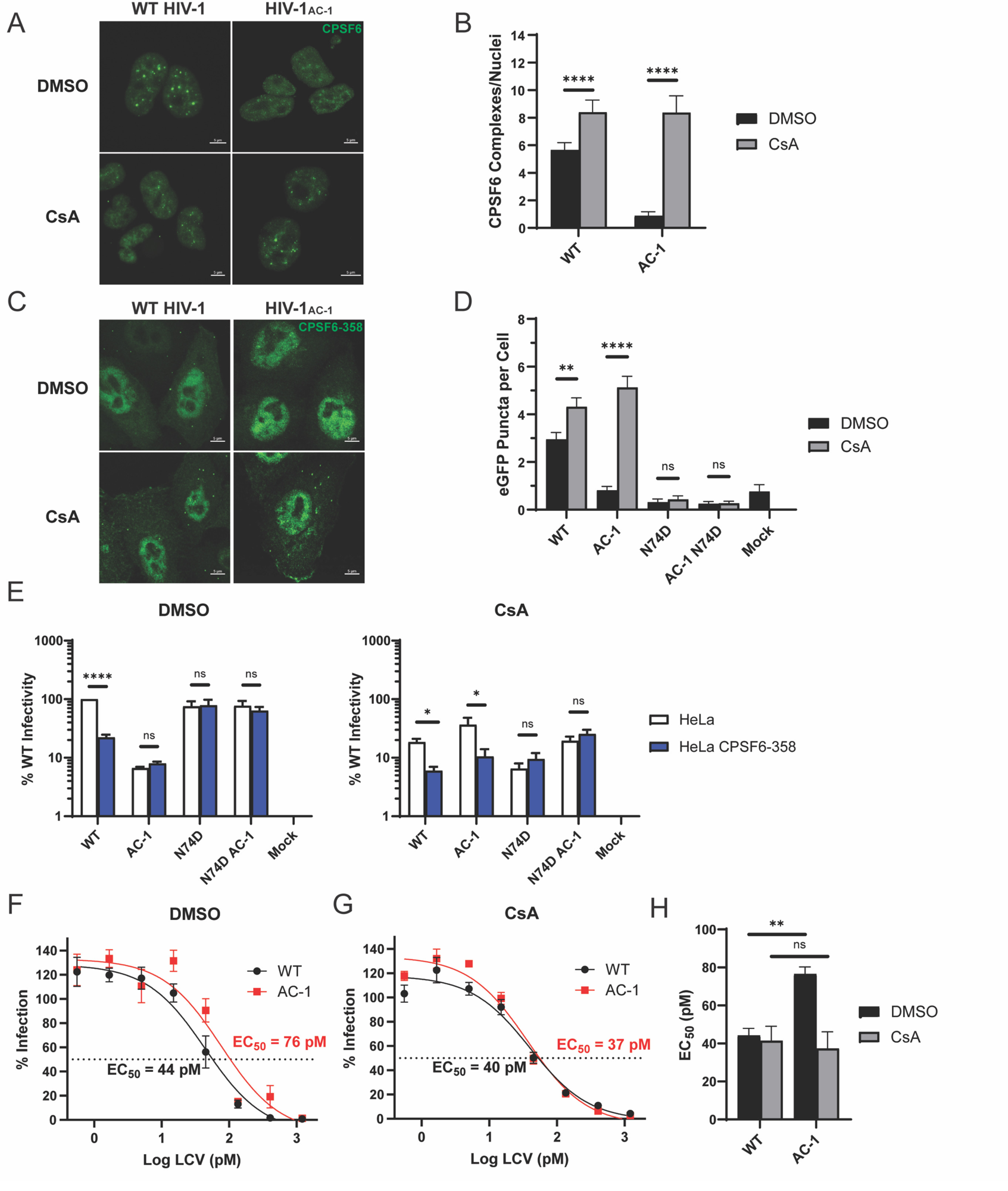
Increased CypA binding to HIV-1 capsid reduces CPSF6 and LCV binding. (A) Representative confocal microscopy images are shown of nuclear CPSF6-GFP higher order complexes in HeLa cells 6 h after WT HIV-1 or HIV-1_AC-1_ infection in DMSO or 10 μM CsA. Scale bars denote 5 μm. (B) CPSF6-GFP higher order complexes shown in A were quantified (n = 3). (C) Representative confocal microscopy images are shown of CPSF6-358-GFP higher order complexes in HeLa cells 30 min after WT HIV-1 or HIV-1_AC-1_ infection in DMSO or 10 μM CsA. Scale bars denote 5 μm. (D) CPSF6-358-GFP higher order complexes shown in C were quantified (n = 3). (E) HeLa cells or HeLa cells expressing CPSF6-358 were infected with WT and mutant HIV-1 in the presence of DMSO or 10 μM CsA (n = 3). Luciferase activity was measured after 48 h. (F-G) LCV sensitivity was measured for WT HIV-1 or HIV-1_AC-1_ in HeLa cells treated with DMSO (F; n = 3) or 5 μM CsA (G; n = 2). (H) The differences in EC_50_ values of the two viruses in the absence (p = 0.0037) or presence (p = 0.76) of CsA are shown. Error bars represent SEM. Comparisons between infection conditions (B, D, E) or virus EC_50_ values (H) were analyzed by unpaired t tests. P values < 0.05 were considered significant and significant values are denoted as *, p < 0.05; **, p < 0.01; ***, p < 0.001; and ****, p < 0.0001. ns, p > 0.05.

Next, we assessed whether increased CypA binding affinity prevents cytoplasmic CPSF6 binding. Truncated CPSF6-358 lacks the RS-like domain required for TNPO3 binding, leading to cytoplasmic accumulation while maintaining HIV-1 capsid binding (Lee et al., 2010). CPSF6-358-GFP forms visible higher order complexes in the cytoplasm upon binding to HIV-1 capsid (Ning et al., 2018). HeLa cells expressing CPSF6-358-GFP were synchronously infected for 30 min with or without CsA treatment prior to fixation and imaging. CPSF6-358-GFP higher order complexes were quantified by excluding the nuclei of cells during analysis and detecting GFP puncta above background in the cytoplasm. As we have previously shown, WT HIV-1 produced visible CPSF6-358-GFP complexes in cells, which increased in the presence of CsA (**Figures 5C, 5D**). N74D HIV-1 and N74D HIV-1_AC-1_, which do not bind CPSF6, failed to produce CPSF6-358-GFP complexes, as expected. HIV-1_AC-1_ failed to form CPSF6-358-GFP higher order complexes in DMSO (**Figures 5C, 5D**). But loss of CypA binding by CsA treatment restored CPSF6-358-GFP complex formation to WT HIV-1 levels, further indicating that CypA binding to HIV-1_AC-1_ capsid prevents cytoplasmic CPSF6 binding.

HIV-1 capsid binding to CPSF6-358 leads to aberrant uncoating, microtubule trafficking and nuclear import, which inhibits infection (Lee et al., 2010; Ning et al., 2018; Zhong et al., 2021). To ascertain whether CPSF6-358 restricts HIV-1_AC-1_, HeLa cells with or without CPSF6-358-GFP expression were infected with mutant viruses (**Figure 5E**). WT HIV-1 infection was significantly restricted in HeLa CPSF6-358-GFP cells compared to control cells, regardless of CsA treatment. In contrast, HIV-1_AC-1_ infection was not restricted by CPSF6-358 unless CsA was present, again indicating that CPSF6-358 binding to capsid was inhibited by CypA. Both N74D HIV-1 and N74D HIV-1_AC-1_ remained insensitive to CPSF6-358, as expected given the loss of CPSF6 binding.

Since increased CypA binding results in a loss of CPSF6 interaction with the HIV-1 capsid, we hypothesized that binding of LCV to HIV-1_AC-1_ also would be reduced compared to WT HIV-1. LCV binds in the hydrophobic pocket targeted by CPSF6 (Bester et al., 2020). Sensitivity of WT HIV-1 and HIV-1_AC-1_ to LCV was measured in increasing concentrations with or without addition of CsA (**Figures 5F** and **5G**, respectively). WT HIV-1 was similarly inhibited by LCV regardless of the addition of CsA (**Figure 5H**). In contrast, the LCV 50% effective concentration (EC_50_) to inhibit HIV-1_AC-1_ infection was approximately 2-fold higher than WT HIV-1 in the absence of CsA (**Figure 5H**). However, CsA treatment reduced the EC_50_ for HIV-1_AC-1_ to that of WT HIV-1. This difference is consistent with the difference in CypA binding affinity between WT HIV-1 and HIV-1_AC-1_ (Lahaye et al., 2013).

### Increased CypA binding to HIV-1 capsid leads to mislocalized integration

Following engagement with WT HIV-1 capsid, CPSF6 promotes the localization of viral pre-integration complexes distal from the nuclear envelope to SPADs and gene rich chromatin, resulting in selective integration into these regions (Chin et al., 2015; Francis et al., 2020a; Li et al., 2020; Singh et al., 2022; Sowd et al., 2016). In contrast, in cells lacking CPSF6 or if capsid cannot bind to CPSF6, HIV-1 integration is mislocalized near the nuclear envelope to DNA containing lamina-associated domains (LADs). Because CPSF6 binding to HIV-1_AC-1_ capsid is reduced due to higher CypA affinity, we assessed whether HIV-1_AC-1_ integrates away from SPADs and gene-dense regions. HeLa cells were infected with HIV-1_AC-1_ or WT HIV-1 for 72 h in DMSO or CsA. HIV-1 proviral integration sites were amplified, sequenced, and analyzed for association with genes, gene density (genes per Mb), SPADs, and LADs, as previously described (Serrao et al., 2016)(Li et al., 2020). N74D HIV-1 and N74D HIV-1_AC-1_ were included as controls that do not bind to CPSF6 and P90A HIV-1 was included as a control that does not bind to CypA. All viruses integrated into genes above the calculated random integration control (RIC) value **(Figure 6A)**. WT HIV-1 and P90A HIV-1 integration sites were highly enriched in gene-dense regions (>20 genes/Mb) and with SPADs (**Figures 6B, 6C**), while avoiding LAD-associated DNA (**Figure 6D**). Notably, treatment with CsA led to significant increases in WT HIV-1 integration in genes, gene dense regions, and SPADs, which is consistent with a previous report (Schaller et al., 2011) and may be due to increased CPSF6 binding. As expected, N74D HIV-1 and N74D HIV-1_AC-1_, which do not bind to CPSF6, disfavored gene-dense regions and SPADs, with concomitant increases in LAD-tropic targeting, regardless of CsA treatment.

**Figure 6:**
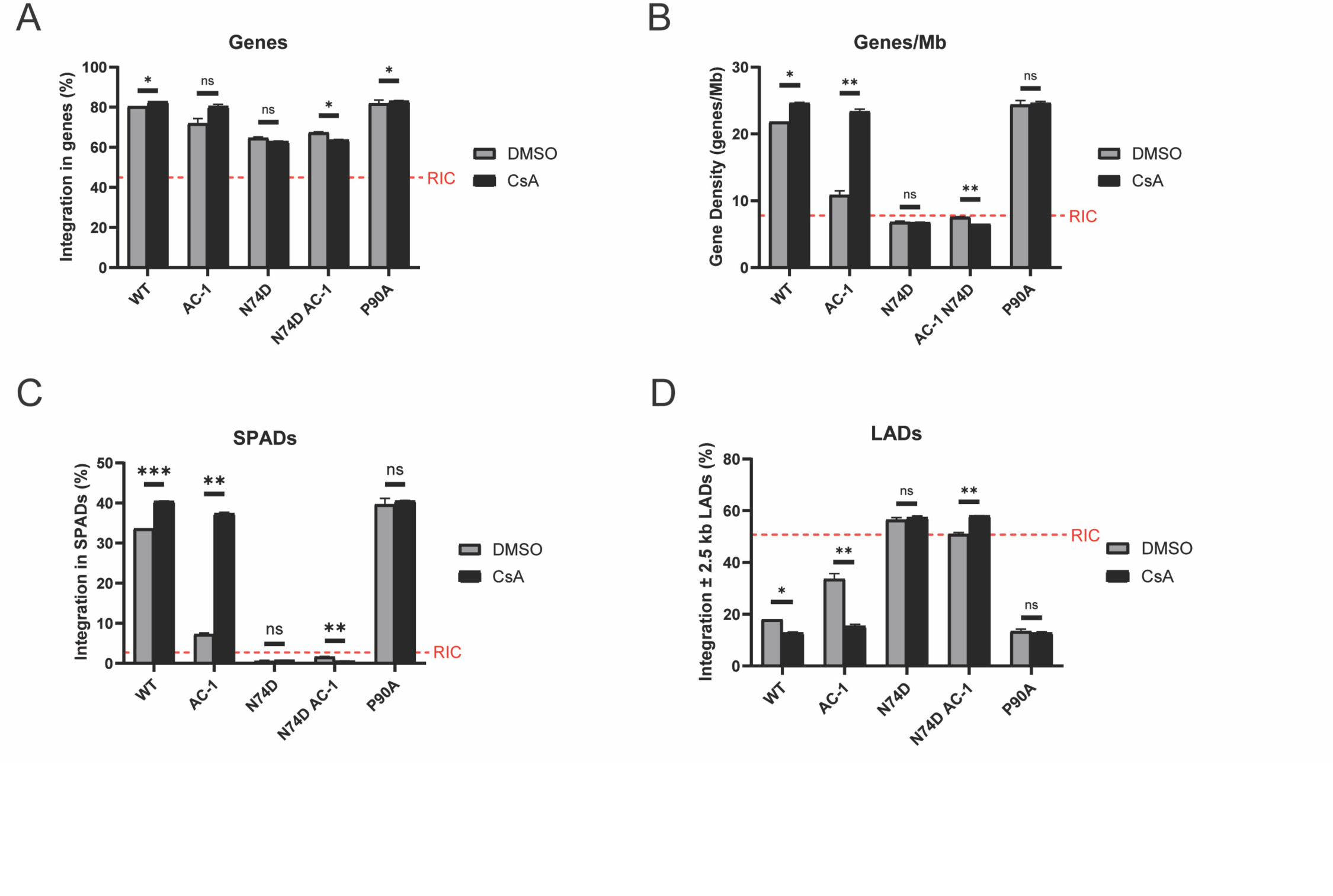
Increased CypA binding to HIV-1 capsid leads to mislocalized integration. WT and mutant HIV-1 integration sites in HeLa cells treated with DMSO or 10 μM CsA were analyzed for genes (A), gene density (B), in SPADs (C), and ± 2.5 kb of LADs (D). Random integration control (RIC) values (red dashed lines) were calculated computationally via mapping 112,183 random integration sites onto human genome build 19 in silico following in silico DNA shearing (Li et al., 2020). The infection and integration site sequencing assays were performed twice and error bars represent SEM. Comparisons between infection conditions were analyzed by unpaired t tests. P values < 0.05 were considered significant and significant values are denoted as *, p < 0.05; **, p < 0.01; ***, p < 0.001; and ****, p < 0.0001. ns, p > 0.05.

Consistent with lower CPSF6 binding, HIV-1_AC-1_ infection in DMSO led to an intermediate integration pattern in DNA with an average gene density of ∼10 genes/Mb and greater affinity for integration into LADs compared to WT HIV-1 (**Figures 6B, 6D**). In contrast, CsA treatment of cells infected with HIV-1_AC-1_ rescued integration to a phenotype similar to WT HIV-1. These results suggest the importance of both CypA and CPSF6 binding to capsid for optimal proviral integration.

## Discussion

Previously, the HIV-1_AC-1_ mutant was created to increase CypA binding affinity to capsid to study immune sensing of HIV-1 infection in myeloid cells (Lahaye et al., 2013). Lahaye and colleagues determined that this mutant had increased CypA binding (approximately 2-fold), which significantly inhibited HIV-1 infection at the nuclear import step, as measured by 2-LTR circles, in all cell types tested (Lahaye et al., 2016). By measuring 2-LTR circles and in a nuclear import kinetics assay, we also showed that HIV-1_AC-1_ capsid translocation through the NPC is significantly reduced. The infectivity defect was abrogated by either treatment of cells with an inhibitor of CypA binding to CA (CsA), knockout of CypA in cells (*PPIA^-/-^*cells), or addition of a CA amino acid substitution to prevent CypA binding (P90A), demonstrating that CypA binding to capsid directly mediates nuclear import and promotes replication.

Additionally, introduction of a mutation that prevents CPSF6 binding to capsid, N74D, rescued HIV-1_AC-1_ infectivity, suggesting that the phenotype is caused by capsid binding to CPSF6. Our previous studies showed that CPSF6 binding to HIV-1 capsid mediated cytoplasmic trafficking on microtubules and uncoating (Ning et al., 2018; Zhong et al., 2021). HIV-1 nuclear import is enhanced by CPSF6 binding to TNPO3 for nuclear localization (Chin et al., 2015; De Iaco et al., 2013; Fricke et al., 2013; Lee et al., 2010). Furthermore, CypA binding to HIV-1 capsid shielded it from binding to cytoplasmic CPSF6 (Ning et al., 2018; Zhong et al., 2021) and TRIM5α (Kim et al., 2019; Selyutina et al., 2020). And while localization of CPSF6 was predominantly nuclear with some higher order complexes detected in the cytoplasm near the nucleus and associated with microtubules, CypA in cell lines, including CD4+ T cells (Zhong et al., 2021), localized to the cell periphery and was excluded from regions of CPSF6 expression. With increased CypA binding to HIV-1_AC-1_ capsid, the interaction with CPSF6 is compromised in the cytoplasm and the nucleus. Furthermore, relocalization of CypA to only the nucleus led to infection phenotypes of WT HIV-1 and HIV-1_AC-1_ similar to those in cells lacking CypA, Thus, our model of optimal HIV-1 replication suggests that abundant CypA binds to the incoming capsid at the periphery of the cell and the availability of CypA decreases near the nucleus, allowing CPSF6 to engage with the capsid, leading to a competitive exchange of capsid binding host factors that is spatiotemporally regulated (**Figure 7**). Similarly, HIV-1_AC-1_ sensitivity to LCV was reduced, suggesting that CA inhibitor binding was diminished in the presence of greater CypA binding.

**Figure 7:**
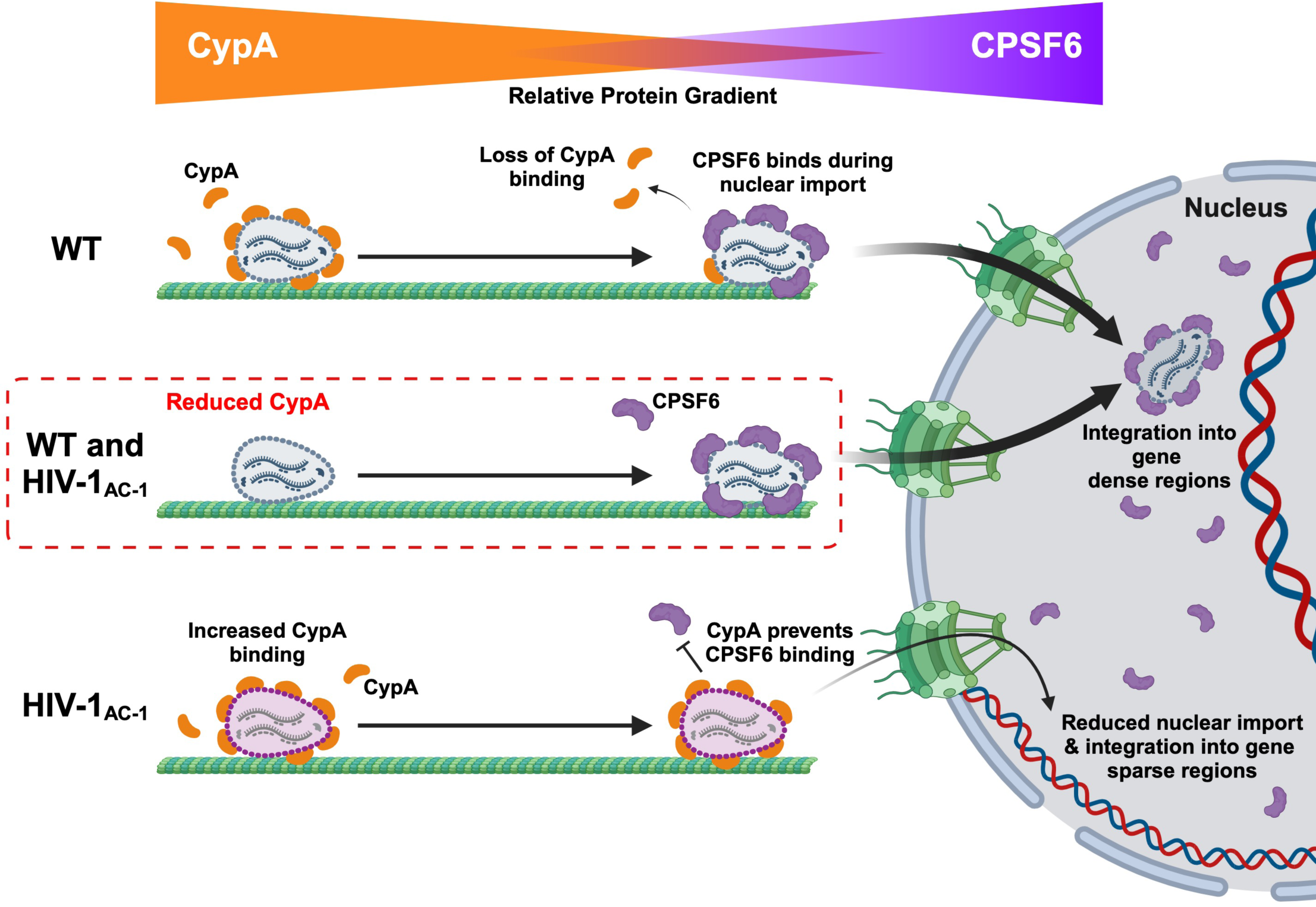
Schematic model of HIV-1_AC-1_ replication and restriction. Cytoplasmic WT HIV-1 engages with CypA to sterically block premature CPSF6 binding. As the HIV-1 capsid approaches the nucleus, CypA concentration decreases, leading to an exchange of CypA for CPSF6 binding in preparation for nuclear import. HIV-1_AC-1_ displays increased CypA binding that results in failed nuclear import due to inability to engage cytoplasmic CPSF6.

At the nucleus, HIV-1 capsid acts like a nuclear transport receptor, binding to phenylalanine-glycine (FG) repeats of nucleoporin condensates to enter the nucleus via NPCs (Dickson et al., 2024; Fu et al., 2024). CPSF6 binding to capsid-like particles competes with nucleoporin FG binding (Dickson et al., 2024), likely in a similar manner of sequential host factor binding to HIV-1 capsid. Previous studies proposed that while CPSF6 is not required for entry through the NPC, it is needed for capsid uncoating and migration away from the inner NPC within the nucleus (Chin et al., 2015; Zila et al., 2019). CypA binding to capsid is required for interaction with inner NPC proteins POM121 and Nup35 (Xue et al., 2023), further suggesting a direct role of CypA in nuclear entry. One can envision a precise sequence of host factors binding to HIV-1 capsid to move it from the periphery of the cell to the cytoplasmic face of the NPC and then through the NPC. However, in the case of HIV-1_AC-1_, nuclear import through the NPC may be further hindered due to excessive CypA binding. Investigation of this mutant with different NUP-FG condensates with and without CypA would provide additional information on the role of CypA in shielding specific NPC components.

After entry into the nucleus, HIV-1 capsid interacts with the CPSF6 FG repeats/prion-like domain (Guedan et al., 2021; Price et al., 2012; Wei et al., 2022) to enable the viral replication complexes to be trafficked further into the nucleus for integration into gene-rich SPADs (Chin et al., 2015; Francis et al., 2020b; Sowd et al., 2016). Surprisingly, HIV-1_AC-1_ had similar *in vitro* capsid stability as WT HIV-1. This was validated in cells by the introduction of CsA at different time points, which led to rescue of HIV-1_AC-1_ infectivity up to 6 h post-infection, indicating that the HIV-1_AC-1_ capsid is likely largely intact. In addition, HIV-1_AC-1_ completed reverse transcription, which has been shown to occur even in hyperstable capsids during infection (Guedan et al., 2021). Beyond 6 h, HIV-1_AC-1_ infectivity could not be rescued, suggesting that an irreversible change occurred in the capsid that could not be overcome even upon the loss of CypA binding. Any HIV-1_AC-1_ capsids that entered the nucleus, perhaps during mitosis, appeared not to engage nuclear CPSF6 in a manner consistent with WT HIV-1, leading to misintegration near the nuclear membrane in LADs (**Figure 6**). This is similar to N74D HIV-1 that is unable to bind to CPSF6 (Li et al., 2020; Sowd et al., 2016). Interestingly, the mislocalization of HIV-1_AC-1_ integration was not as pronounced as N74D HIV-1, suggesting that some CPSF6 binding can occur in the nucleus. In contrast, with a loss of CypA binding, HIV-1_AC-1_ integrated preferentially into gene-dense regions and SPADs while avoiding gene sparse LADs, similar to WT HIV-1.

Our factor exchange model focused on the spatial and temporal aspects of host factors binding to HIV-1 capsid, which could be influenced by amino acid substitutions. Surprisingly, the introduction of the five CA amino acid substitutions to create HIV-1_AC-1_ results in a relatively small change in binding affinity to CypA (Lahaye et al., 2013) that significantly impacts viral replication. This suggests that a weaker binding affinity of CypA to HIV-1 capsid allows a proper host factor exchange with CPSF6 for optimal nuclear import and integration. Analysis of over 20,000 HIV-1 group M Gag sequences show that five of the six HIV-1_AC-1_ substitutions are highly polymorphic (Tao et al., 2023). Proline is the consensus residue at position 92 and Ile86, Leu91, and Leu96, all HIV-1_AC-1_ changes, are observed at up to 10% prevalence. The sixth substitution, proline to alanine at position 93, has a frequency of < 1% in these sequences. It is possible that CypA binding to HIV-1 capsid may be variable in people living with HIV, which may lead to aberrant host factor binding during replication and could affect pathogenesis. Furthermore, some HIV-1 variants similar to HIV-1_AC-1_ may have reduced sensitivity to LCV treatment, which may limit its efficacy and promote development of resistance mutations in some people. Our model incorporates spatiotemporal regulation of host protein composition on capsids that promotes or restricts HIV-1 replication. Importantly, this model can be expanded beyond CypA and CPSF6 to other cell factors that interact with capsid during infection.

## Materials and Methods

### Cell lines and primary cultures

HeLa, HeLa cells stably expressing CPSF6-358-eGFP (Ning et al., 2018), HEK293T, HT1080, and HT1080 CypA^-^/^-^ cells (Layish et al., 2024) were grown in culture with Dulbecco’s Modified Eagle Medium (DMEM; Thermo Fischer Scientific) supplemented with 10% fetal bovine serum (FBS; Atlanta Biologicals) and 100 μg/ml penicillin, 100 μg/ml streptomycin, and 2 mM L-glutamine (PSG; Thermo Fischer Scientific) at 37° C and 5% CO_2_. Jurkat T cells and Jurkat *PPIA*^-/-^ cells (obtained through the NIH HIV Reagent Program, Division of AIDS, NIAID, NIH; ARP-10095, contributed by Drs. D. Braaten and J. Luban) were cultured in RPMI medium supplemented with 10% FBS, PSG at 37° C and 5% CO_2_. Primary CD4+ T cells were isolated from whole blood (BioIVT) using the CD4+ T Cell Isolation Kit (Miltenyi Biotec) and were cultured in RPMI medium containing 10% FBS, PSG, and 20 IU/ml IL-2 and stimulated with 2 μg/mL PHA for 48 h prior to infection. GHOST-R3/X4/R5 cells (Cecilia et al., 1998) were maintained in DMEM medium containing 10% FBS, PSG, 1 μg/ml puromycin, 100 μg/ml hygromycin, and 500 μg/ml geneticin at 37° C and 5% CO_2_.

### Plasmids and viruses

Viruses were produced by transfection of HEK293T cells using either Lipofectamine 2000 (Invitrogen) or polyethylenimine (PEI; Polysciences). The plasmids encoding replication-defective HIV-1 with firefly luciferase or mNeonGreen in place of *nef* (pNLdE-luc and pNLdE-mNeon, respectively) and pL-VSV-G were used to produce pseudotyped HIV-1 (Lee et al., 2010). The HIV-1_AC-1_ mutations were introduced into WT and N74D pNLdE-luc using the Gibson Assembly Cloning kit (New England Biolabs) and primers shown in **Table S1**. The P90A substitution was generated via the QuikChange Mutagenesis Kit using primers shown in **Table S1**. Fluorescently labeled IN was packaged into viruses using the pVpr-mRuby3-IN and pVpr-mNeon-IN plasmids during transfection (Ning et al., 2018). Labelling of viruses was confirmed by imaging viruses in MatTek dishes by total internal reflection fluorescence (TIRF) microscopy. Lentiviral vectors encoding control or TRIM5_α_ miRNAs (pAPM-D4-miR30-L1221 or pAPM-D4-miR30-TRIM5_α_; Addgene) were produced by co-transfection with the packaging plasmid pcHelp (Mochizuki et al., 1998) and pCMV-VSV-G (Kim et al., 2019). All viruses were harvested by passing cell supernatants through a 0.45 mm filter before storing at -80° C. HIV-1 titers were determined in GHOST-R3/X4/R5 cells and total virus was quantified by p24 enzyme-linked immunosorbent assay (ELISA) (XpressBio). N-tropic MLV was produced by transfection of the pCIG3 N packaging plasmid (a gift from Jonathan Stoye) (Bock et al., 2000), the pFB-Luc vector (Agilent), and pCMV-VSV-G.

The plasmid pLVX-hNUP62Fv2GFP for the NIK assay was a gift from Edward Campbell (Dharan et al., 2020). Plasmids pLBCX-GS-HA-eGFP-CPSF6 and pLBCX-GS-HA-eGFP-CPSF6/F284A to express GFP fusions of the 551-residue isoform of CPSF6 (WT or F284A) were constructed in plasmid vector pLB(N)CX essentially as previously described (Sowd et al., 2016). CypA was amplified by PCR from cDNA produced with random hexamers from RNA isolated from HeLa cells. The PCR product was Topo cloned into the plasmid pcDNA3.1 (Thermo Fisher) to produce pCypA. The SV40 NLS was added to the C-terminus of CypA via annealing partially complimentary primers encoding the SV40 NLS (**Table S1**) at 20 μM with NEB buffer 2 (New England Biolabs) at 95° C for 4 min and 70° C for 10 min, followed by cooling to room temperature over 4 h. The annealed primers were gel-purified and formed overhangs consistent with BamHI and NotI restriction digests and cloned into pCypA-DsRed (Francis et al., 2016), which had been digested with BamHI and NotI, to replace DsRed with the NLS, producing pCypA-NLS.

### HIV-1 infectivity assays

HeLa cells (5 x 10^4^) were infected in technical duplicates with VSV-G pseudotyped HIV-1 (10 ng p24) in the presence of 5-10 μM CsA in DMSO or equivalent concentrations of DMSO. For growth arrest, cells were treated for 18 h with media containing 1 μg/ml aphidicolin. For knockdown of TRIM5_α_, cells were transduced with lentiviral vectors expressing control or TRIM5_α_ miRNA for 48 h followed by 48 h of selection in media containing 2 μg/ml puromycin prior to infection with HIV-1 in the presence or absence of CsA. For controls for aphidicolin and miRNA treatment, cells were also infected with N-MLV. Cells were either lysed or fixed with 2% paraformaldehyde 48 h later. Infectivity was measured either by luciferase using the Luciferase Assay System Kit (Promega) or by flow cytometry. HT1080 and HT1080 CypA^-^/^-^ cells were transfected with pcDNA3.1, pCypA, or pCypA-NLS 48 h prior to HIV-1 infection in the presence or absence of CsA.

### CA retention assay

The CA retention assay was performed on WT HIV-1 and HIV-1_AC-1_, using mutant viruses with CA E45A or K203A as controls, as previously described (Xu et al., 2020). Briefly, chamber slides (Greiner Bio-One) were treated with Cell-Tak (Corning) for 1 h prior to adherence of viruses. HIV-1 (5 ng p24) containing mRuby3-IN was added to each well in STE buffer (100 mM NaCl, 10 mM Tris-HCl, 1 mM EDTA, pH 7.4). Samples were mounted in gelvatol and imaged by TIRF on a Nikon Eclipse Ti microscope. Six fields per condition were imaged and CA and mRuby3-IN signals were enumerated using Nikon Elements software.

### Imaging of CPSF6 and CypA

HeLa cells stably expressing CPSF6-358-eGFP were infected in the presence or absence of CsA and imaged as previously described (Ning et al., 2018). To detect full-length CPSF6 in cells, HeLa cells were transfected with pLBCX-GS-HA-eGFP-CPSF6 or pLBCX-GS-HA-F284A eGFP-CPSF6 plasmids 48 h prior to replating and infection with HIV-1. Synchronized HIV-1 infections were performed by adding virus in media at 4°C for 15 min to allow cell attachment. Media was removed and replaced with 37°C media to initiate synchronized fusion. After 6 h, cells were fixed in 2% paraformaldehyde. Fixed samples were permeabilized with 0.1% Triton-X100 before staining with Hoechst 33342. Imaging was performed on a Nikon A1 confocal microscope with a motorized piezo Z stage. Z-stacks were created before enumerating nuclei and CPSF6 higher order complexes in 12 fields of view using Nikon Elements.

HT1080 and HT1080 CypA^-/-^ cells with or without transfection with pcDNA3.1, pCypA, or pCypA-NLS were fixed with 2% PFA. Cells were permeabilized with 0.1% Triton-X100 before staining with mouse anti-CypA (Abcam AB58144; 0.48 μg/ml) and rabbit anti-CPSF6 (Novus NBP1-85676; 0.75 μg/ml) antibodies. Samples were then stained with either donkey anti-mouse-Cy3 (Jackson Immuno Research 715-165-151; 1:1000) or donkey anti-rabbit-AlexaFluor488 (Thermo Fisher A-21206; 1:750), respectively. Samples were additionally stained with Hoescht 33342 before mounting samples and imaging individual Z-slices on a Nikon A1 confocal microscope with a motorized piezo Z stage.

### NIK assay

HeLa cells were transfected with pLVX-hNUP62Fv2GFP and selected in medium containing 2 μg/ml puromycin. Cells were growth arrested with 1.5 μM aphidicolin 24 h prior to infection. Cells were synchronously infected with equal amounts of p24 in the presence or absence of CsA. At each timepoint, media containing 1.5 μM AP20187, a homodimerizing drug (MedChemExpress), was added to the cells. Media was replaced at 24 h and cells were lysed 24 h later for measurement of luciferase. Infectivity of each time point was normalized to cells infected in the absence of AP20187.

### Measurement of HIV-1 reverse transcripts and 2-LTR circles

HeLa cells treated with or without CsA were infected with HIV-1 (10 ng p24) that had been treated with 100 U/ml of DNase I (Roche) for 1 h at 37° C. Rilpivirine (1 μM; NIH HIV Reagent Program) was added as a control to parallel cultures to prevent reverse transcription and control for plasmid carryover that may persist in virus stocks. After 24 h, the cells were removed with trypsin-EDTA, washed, and pelleted. Genomic DNA was extracted using the QIAmp DNA Mini Kit (Qiagen). Reverse transcript products (*gag* or second strand transfer) and 2-LTR circles were measured by quantitative PCR using primers and probes (**Table S2**), as previously described (Julias et al., 2001).

### Integration site analysis

HeLa cells treated with or without CsA were infected with DNase-treated HIV-1 at a multiplicity of infection (MOI) of 1 and were trypsinized 3 days post-infection. Cell pellets were stored at -20° C. Genomic DNA was extracted from all samples using the QIAmp DNA Mini Kit. Integration libraries were prepared using ligation-mediated PCR (LM-PCR) as described previously (Matreyek et al., 2014; Serrao et al., 2016). Genomic DNA (5 µg) from HIV-1 infected cells was digested and ligated with linkers using the NEBNext Ultra II FS DNA Library Prep Kit (New England Biolabs) and its associated protocol. In short, genomic DNA was digested in a final reaction volume of 35 µl, containing 7 µl NEBNext Ultra II FS reaction buffer and 2 µl of NEBNext Ultra II FS enzyme Mix, at 37° C for 20 min. The FS enzyme mix was inactivated by heating the reaction at 65° C for 30 min. Three independent ligation reactions of a final volume of 68.6 µl were performed, containing 2.5 µl 10 µM double stranded asymmetric linkers with a 3’-T overhang and 30 µl NEBNext Ultra Ligation Master Mix and 1 µl NEBNext Ligation Enhancer at 20° C for 16 h. Purified ligation products were used in nested PCRs to amplify viral-host integration junctions. Purified PCR products were subjected to 150 bp paired-end Illumina sequencing at Genewiz.

Illumina raw reads were processed, integration sites were determined against the hg19 human genome, and analyzed with various genomic features, as per previously described methodologies (Anderson-Daniels et al., 2019; Li et al., 2020). The coordinates of SPADs, LADs, and RIC were as described previously (Achuthan et al., 2018; Francis et al., 2020b; Li et al., 2020).

### LCV sensitivity assays

LCV sensitivity was determined for WT HIV-1 and HIV-1_AC-1_ as previously described (Melody et al., 2015). Briefly, HeLa cells treated with CsA or equivalent concentration of DMSO were infected for 48 h at a MOI of 0.003 in phenol red-free medium containing dilutions of LCV (NIH HIV Reagent Program) in DMSO. Cells with virus only or without virus and drug were used as positive and negative controls, respectively. Luciferase activity was measured using Britelite Plus (PerkinElmer). Luciferase was normalized to the 100% infection control (no drug) and background (no virus and no drug) was subtracted. The effective concentration to inhibit 50% of virus replication (EC_50_) was calculated by log transforming drug concentrations and using a four-parameter variable slope nonlinear regression for curve fitting analysis by Prism.

### Statistical analysis

Statistical analysis for all was performed in Prism 10 (GraphPad) with statistical tests described in the figure legends. Experiments were completed with technical replicates with 2-3 experimental replicates. Statistical p values are shown with the following designations: p > 0.05, ns; *, p < 0.05; **, p < 0.01; ***, p < 0.001; and ****, p < 0.0001.

## Data availability

Illumina raw sequences for integration sites are available at the National Center for Biotechnology Sequences Read Archive with accession number XXXXXX.

## Acknowledgements

The authors thank Chandra Nath Roy, and Ella Begovic for technical assistance. This work was supported by National Institutes of Health (NIH) grants U54 AI170791 (A.N.E., S.C.W., and Z.A.), R01 AI052014 (A.N.E.), and T32 AI049820 (Z.I.) and the Gilead Sciences Research Scholars Award (M.K.).

## Author Contributions

Conceptualization: Z.I. and Z.A.; Methodology: Z.I., P.K.S., H.L.F., and Z.A.; Investigation: Z.I., C.K., A.K.H., and P.K.S.; Formal Analysis: Z.I., P.K.S., and Z.A.; Resources: S.C.W. and M.K.; Writing - Original Draft: Z.I. and Z.A.; Writing - Review & Editing: all authors; Supervision: A.N.E. and Z.A.; Funding Acquisition: Z.I., A.N.E., and Z.A.

