## Supplemental data for "Spatiotemporal binding of cyclophilin A and CPSF6 to capsid regulates HIV-1 nuclear entry and integration"

**Table S1: Primers used for cloning and mutagenesis of HIV-1 and CypA constructs**

| Name | Sequence (5' to 3') | Use |
|---|---|---|
| NL43-CAac-1 F | GGATAGATTGCATCCAATACATGCAGGGCCTCTT CCAGCAGGCCAGTTGAGAGAACCAAGGGG | Gibson Assembly for AC-1 pNLdE-Luc |
| NL43-CAac-1 R | CCCCTTGGTTCTCTCAACTGGCCTGCTGGAAGA GGCCCTGCATGTATTGGATGCAATCTATCC | |
| KanR-NcoI F | CCCCTTGGTTCTCTCAACTGGCCTGCT GGAAGAGGCCCTGCATGTATTGGATGCAATCTATCC | Gibson Assembly for AC-1 pNLdE-Luc |
| KanR-NcoI R | CCACCATGATATTCGGCAAGCAGGCAT CGCCATGGGTCACGACGAGATCCTCGCCGTCG | |
| P90A-ac-1 F | CCAATACATGCAGGGGCTCTTCCAGCAGGCCAG | Site Directed Mutagenesis for P90A AC-1 NLdE-Luc |
| P90A-ac-1 R | CTGGCCTGCTGGAAGAGCCCCTGCATGTATTGG | |
| CypA F | AACCGTGTACTATTAGCCATGG | PCR amplification of CypA from human cDNA |
| CypA R | AGTCAAACTTATTCGAGTTGTCC | |
| BamHI-NLS F | GGATCCACCGGTCGCCACCCCGAAAAAAAAACGCAA AGTGGAAGATCCGTAGC | Cloning of NLS onto CypA |
| NLS-NotI R | GCGGCCGCTACGGATCTTCCACTTTGCGTTTTTTTTTC GGGGTGGCGACCGGTG | |

**Table S2: Primers for quantitative PCR**

| Name | Sequence (5' to 3') |
|---|---|
| gag F | GCCTGGGAGCTCTCTGGCTAA |
| gag R | GCCTTGTGTGTGGTAGATCCA |
| gag Probe | FAM-AAGTAGTGTGTGCCCGTCTGTTGTGTGACTC-TAMRA |
| 2-LTR F | TTCGCAGTTAATCCTGGCCTT |
| 2-LTR R | GCACACAATAGAGGACTGCTATTGTA |
| 2-LTR Probe | FAM-TAGAGACATCAGAAGGCTGTAGACAAA-TAMRA |

**Supplemental Figure Legends**

**Figure S1.** (A) Infectivity of WT and mutant HIV-1 (10 ng p24) were determined after 48 h by luciferase activity in DMSO or 10 μM CsA in HeLa cells ($n = 3$). (B) N-MLV infection was determined after 48 h by luciferase activity in HeLa cells transduced with lentiviruses expressing control or TRIM5α miRNA ($n = 2$). Error bars represent SEM. Comparisons between infection conditions were analyzed by unpaired t tests. (C, D) DMSO or 10 μM CsA was added at different time points to HeLa cells synchronously infected with WT HIV-1 (C) or HIV-$1_{AC-1}$ (D). The assay was performed three times. Error bars represent SEM and unpaired t-tests were performed for comparisons between conditions at each time point. P values $< 0.05$ were considered significant and significant values are denoted as *, $p < 0.05$; **, $p < 0.01$; ***, $p < 0.001$; and ****, $p < 0.0001$. ns, $p > 0.05$.

**Figure S2.** Nuclear import kinetics were determined for (A) WT HIV-1 and (B) HIV-$1_{AC-1}$ infection of HeLa cells in DMSO or 10 μM CsA containing media at different time points ($n = 3$). Infectivity was determined by luciferase activity (relative luciferase units, RLU). Error bars represent SEM.

**Figure S3.** (A) Representative confocal microscopy images are shown of nuclear WT or F284A CPSF6-GFP higher order complexes in HeLa cells 6 h after WT HIV-1 infection. Scale bars denote 5 μm. (B) WT and F284A CPSF6-GFP higher order complexes shown in A were quantified ($n = 3$). Error bars represent SEM. The comparison of WT and F284A CPSF6-GFP puncta was analyzed by an unpaired t test. P values of $< 0.05$ were considered significant and significant values are denoted as *, $p < 0.05$; **, $p < 0.01$; ***, $p < 0.001$; and ****, $p < 0.0001$. ns, $p > 0.05$.

### Figure S1

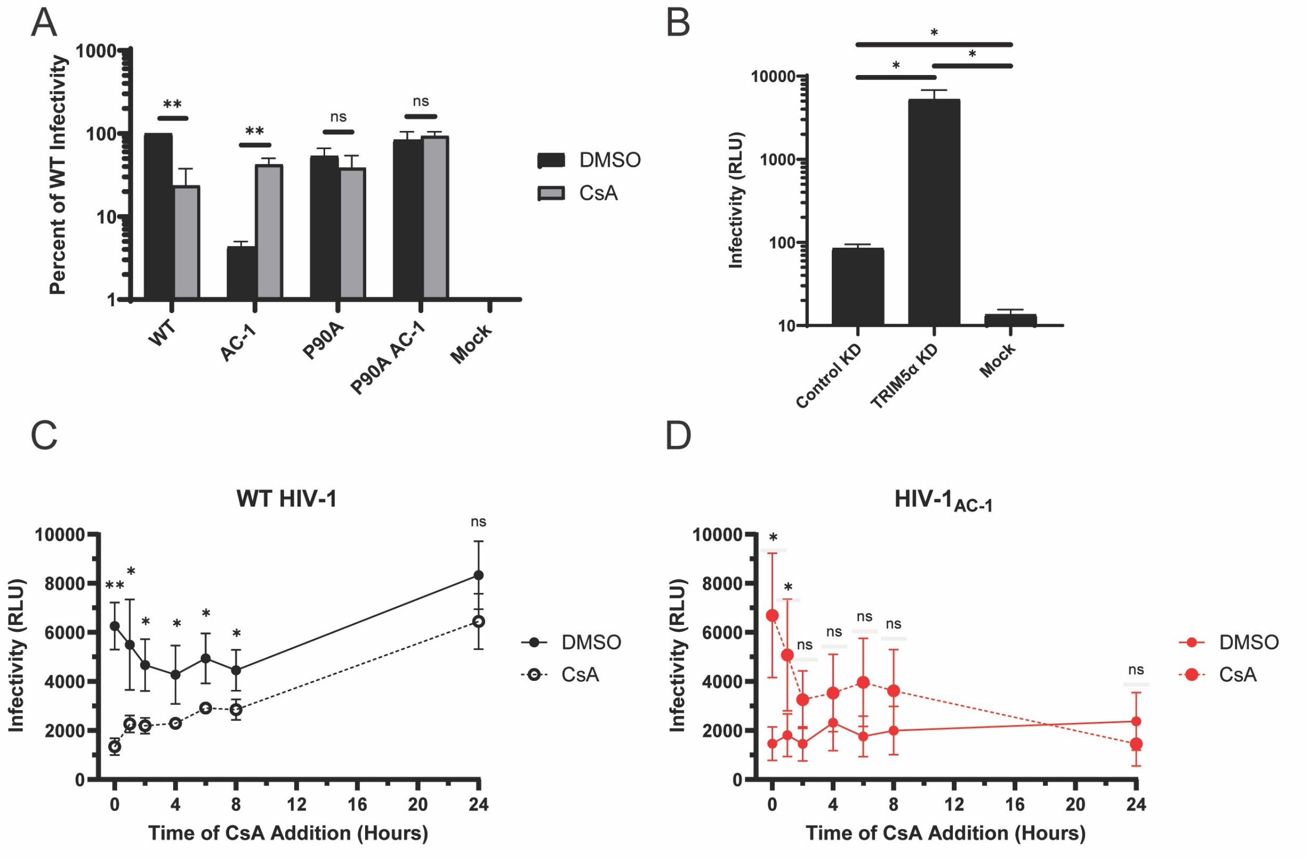
A
B
C
D
Percent of WT Infectivity
1000
100
10
1
**
**
ns
ns
WT
AC-1
P90A
P90A AC-1
Mock
DMSO
CsA
Infectivity (RLU)
10000
1000
100
10
*
*
*
Control KD
TRIM5α KD
Mock
WT HIV-1
HIV-1AC-1
Infectivity (RLU)
10000
8000
6000
4000
2000
0
0
4
8
12
16
20
24
Time of CsA Addition (Hours)
DMSO
CsA
ns

#### Figure S2

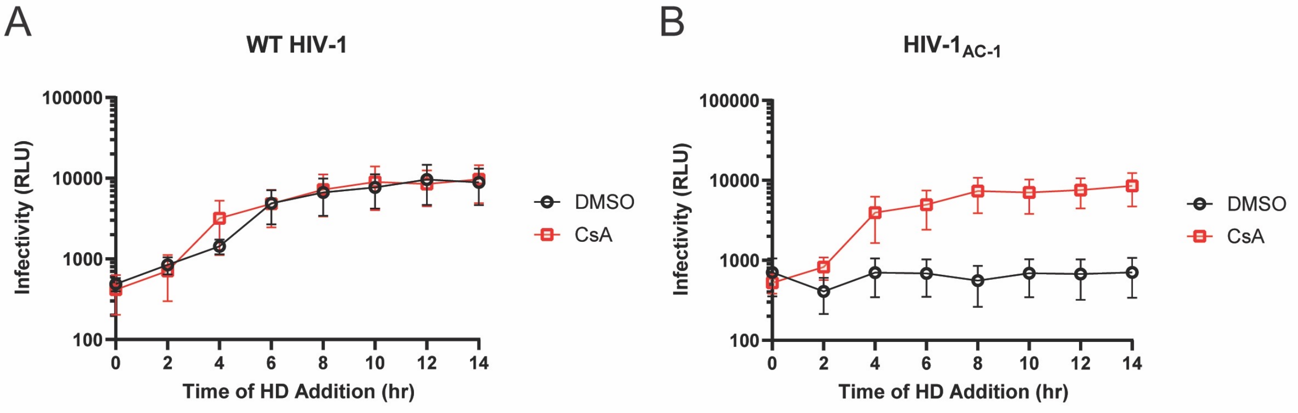
A
WT HIV-1
100000
10000
1000
100
Infectivity (RLU)
0
2
4
6
8
10
12
14
Time of HD Addition (hr)
DMSO
CsA
B
HIV-1$_{AC-1}$
100000
10000
1000
100
Infectivity (RLU)
0
2
4
6
8
10
12
14
Time of HD Addition (hr)
DMSO
CsA

### Figure S3

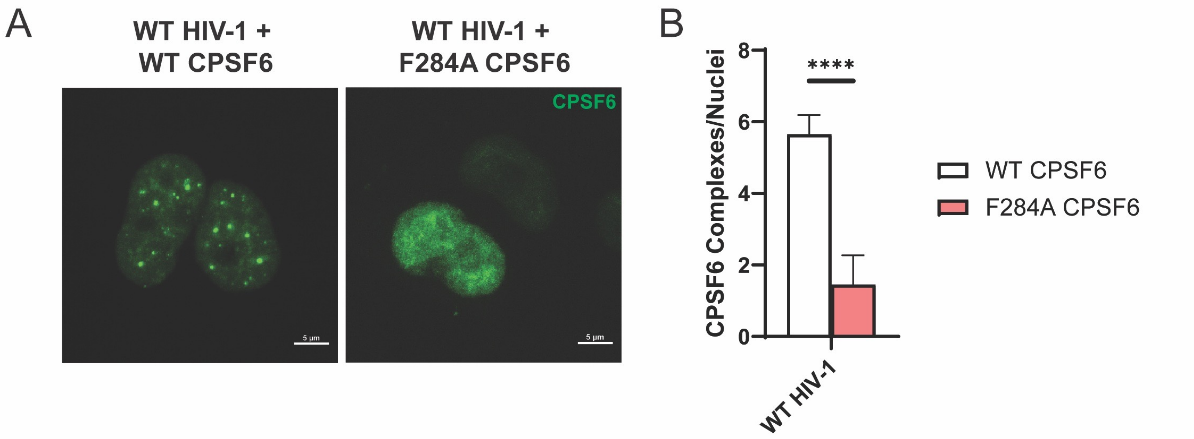
A
WT HIV-1 +
WT CPSF6
WT HIV-1 +
F284A CPSF6
CPSF6
5 μm
5 μm
B
CPSF6 Complexes/Nuclei
8
6
4
2
0
****
WT HIV-1
WT CPSF6
F284A CPSF6
